# Anatomical meniscus construct with zone specific biochemical composition and structural organization

**DOI:** 10.1101/665067

**Authors:** G. Bahcecioglu, B. Bilgen, N. Hasirci, V. Hasirci

## Abstract

A PCL/hydrogel construct that would mimic the structural organization, biochemistry and anatomy of meniscus was engineered. The compressive (380 ± 40 kPa) and tensile modulus (18.2 ± 0.9 MPa) of the PCL scaffolds were increased significantly when constructs were printed with a shifted design and circumferential strands mimicking the collagen organization in native tissue (p<0.05). Presence of circumferentially aligned PCL strands also led to elongation and alignment of the human fibrochondrocytes. Gene expression of the cells in agarose (Ag), gelatin methacrylate (GelMA), and GelMA-Ag hydrogels was significantly higher than that of cells on the PCL scaffolds after a 21-day culture. GelMA exhibited the highest level of collagen type I (*COL1A2)* mRNA expression, while GelMA-Ag exhibited the highest level of aggrecan (*AGG)* expression (p<0.001, compared to PCL). GelMA and GelMA-Ag exhibited a high level of collagen type II (*COL2A1*) expression (p<0.05, compared to PCL). Anatomical scaffolds with circumferential PCL strands were impregnated with cell-loaded GelMA in the periphery and GelMA-Ag in the inner region. GelMA and GelMA-Ag hydrogels enhanced the production of COL 1 and COL 2 proteins after a 6-week culture (p<0.05). COL 1 expression increased gradually towards the outer periphery, while COL 2 expression decreased. We were thus able to engineer an anatomical meniscus with a cartilage-like inner region and fibrocartilage-like outer region.

## 1 Introduction

Meniscus is a semilunar and wedge-shaped fibrocartilage located between the femur and the tibia. The outer periphery of the meniscus is mainly fibrous and contains high amount of type I collagen (COL 1) (70-80% of the total collagen), while the inner region is cartilaginous and contains a high amount of type II collagen (COL 2) (50-60% of the collagen) [1,2]. These collagen fibers are mainly circumferentially oriented and contribute to the tensile properties of the tissue [3,4]. Proteoglycans, which are composed of glycosaminoglaycans bound to a core protein such as aggrecan, decorin and biglycan, are also more abundant in the inner region of the meniscus compared to the outer region, and contribute to the compressive properties of the tissue [2,5,6].

It is a challenge to design an artificial meniscus due to its complex structure. An engineered meniscus should mimic the structural organization, biochemistry and anatomy of the native tissue. Attempts to engineer total meniscal constructs include lyophilized foams [7–9] which are suitable for use in partial replacement of the meniscus due to their low mechanical properties, electrospun mats [10,11] which are too thin to be considered for total meniscal replacement, and combination of the two which still have low mechanical properties [12,13] and some lead to degeneration of the hyaline cartilage [14].

Three dimensional (3D) printing is an additive manufacturing technique that enables production of constructs in the desired shapes and architecture and with appropriate mechanical properties, and thereby is an attractive method to create scaffolds mimicking the meniscus. Alginate [15], collagen [16] and gelatin methacrylate (GelMA) [17] hydrogels have been printed as meniscal constructs, but with poor mechanical properties. Others involve the use of poly(ε-caprolactone) (PCL) [18–22], which has superior mechanical properties. Some of these PCL-based 3D printed scaffolds even include circumferentially oriented strands mimicking the collagen organization in the meniscus [20–22]. However, one major drawback of PCL is that it lacks biofunctional groups that help in cell adhesion and extracellular matrix (ECM) production. This deficiency reduces the amount of ECM produced on these scaffolds [2], as well as limiting the production of region-specific ECM components. In order to mimic the biochemical composition of the meniscus, Lee and colleagues [20] have tethered microspheres carrying transforming growth factor (TGF)-ß1 and connective tissue growth factor (CTGF) on the inner and outer regions of the 3D printed PCL scaffolds, respectively. They have succeeded in engineering a construct with a cartilaginous inner portion and a fibrous periphery. However, PCL scaffolds have been shown to degenerate the underlying articular cartilage mainly because of friction between the stiff polymeric material and the softer natural tissue [20,21].

Hydrogels absorb water and build hydrostatic pressure against compressive load, and thus act as a cushion between the scaffolds and the cartilage and protect them from degeneration [24,25]. Recently, we have shown that agarose (Ag) and GelMA hydrogels protect the cells from the mechanical damage when cells on 3D printed PCL scaffolds were dynamically compressed [26]. Our group, as well as others, have also shown that Ag is chondrogenic and GelMA is fibrogenic [2,23,26]. These findings enable production of zone-specific tissues in the different regions of the meniscal constructs.

In our previous study we impregnated the meniscus shaped PCL scaffolds with porcine fibrochondrocyte-loaded Ag in the inner region and GelMA in the outer region to mimic the biochemical composition of the meniscus [26]. Here, we propose an improved version of our construct. We printed the meniscus shaped PCL scaffolds (i) with shifted strands to decrease the compressive modulus of the constructs and thus reduce the risk of cartilage degeneration after possible implantation and (ii) with circumferentially oriented strands resembling those of the native meniscus to increase the tensile properties, (iii) impregnated the inner region of our constructs with cell-loaded GelMA-Ag instead of Ag to enhance the integrity of the outer and inner regions as well as enhancing the collagen production, and (iv) produced anatomical constructs using the magnetic resonance imaging (MRI) data of a miniature pig to improve the biomechanical performance of the constructs. We also used human fibrochondrocytes to evaluate the performance of our constructs before a possible clinical application. To the best of our knowledge, this construct is the closest scaffold to the native meniscus in terms of anatomy, structure and biochemical composition and could be a promising substitute for use in total meniscus replacement.

## 2 Materials and Methods

### 2.1 Preparation of PCL scaffolds

Square prism (10 mm × 10 mm × 3 mm), rectangular prism (30 mm × 10 mm × 3 mm), and Coliseum (outer diameter: 26 mm, inner diameter: 8 mm, and height at periphery: 5 mm) shaped models were designed using SketchUp software (Google inc., USA) and converted to stereolithography (STL) file formats as described previously [2,26]. The models were transferred to Bioscaffolder (SYS+ENG, SalzgitterBad, Germany) through computer-aided manufacturing (CAM) software (PrimCAM, Einsiedeln, Switzerland). Poly(ε-caprolactone) (PCL, Mw: 50 kDa, Polysciences, USA) scaffolds were printed with strand orientation of 0/90°, strand distance of 1 mm, with or without shifting (offset: 0.5 mm), and with or without circumferential strands (contours) (Fig. 1A(i)). At the end, scaffolds without circumferential strands and having non-shifted (NS) or shifted (S) designs, and scaffolds with circumferential strands and having non-shifted (NSC) or shifted (SC) designs were obtained.

**Figure 1.**
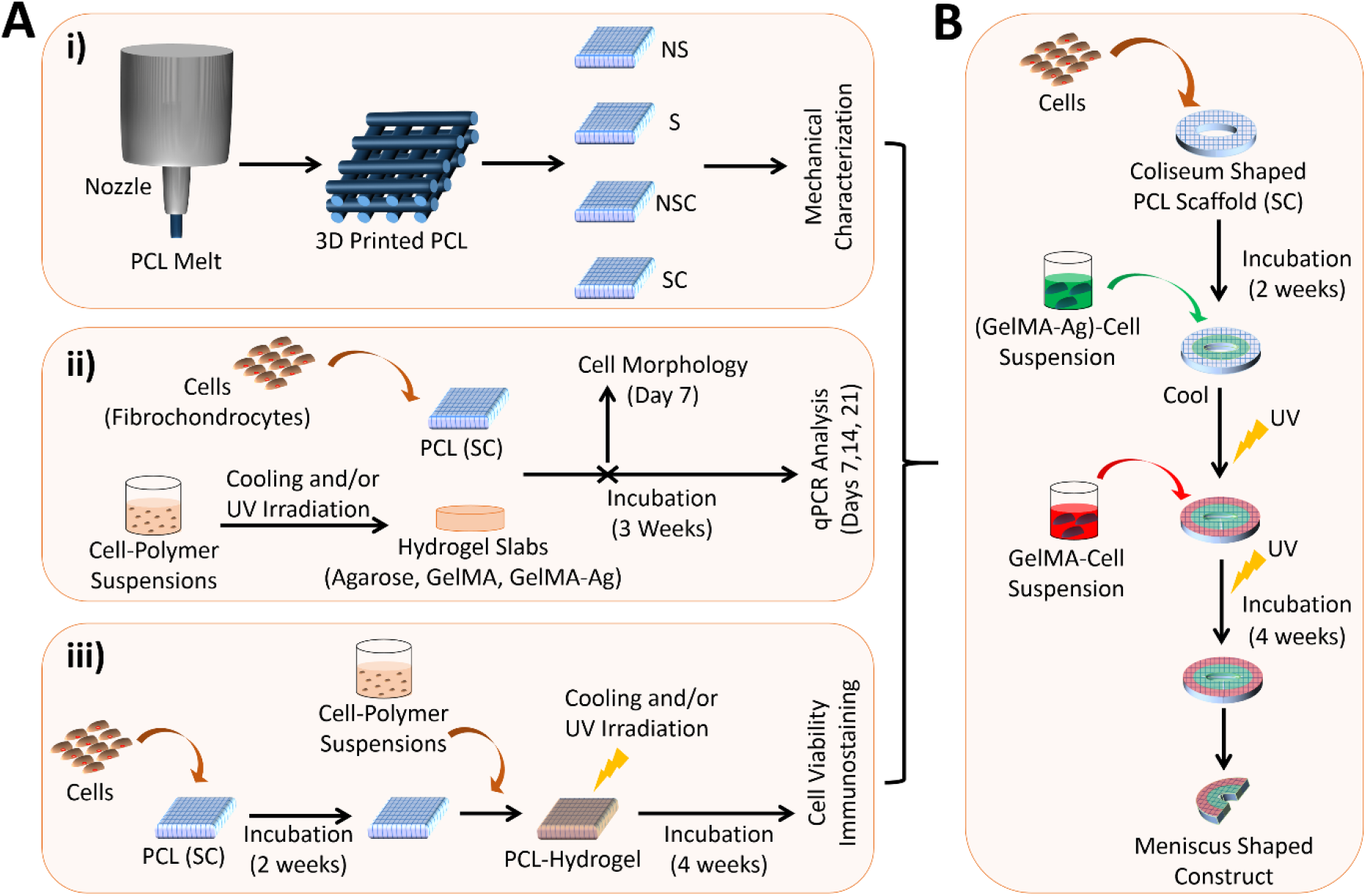
Diagram showing the study design. (A) Optimization of the construct preparation conditions. (i) PCL scaffolds were printed with 0-90° strand orientation, with or without shifting at the consecutive layers and with or without circumferential strands, and characterized mechanically. (ii) 3D printed PCL and slabs of Ag, GelMA, and GelMA-Ag hydrogels were seeded with fibrochondrocytes, incubated for 21 days in culture media, and tested for cell morphology at Day 7, and for mRNA expression at Days 7, 14, and 21. (iii) Square prism PCL scaffolds were incubated for 2 weeks, impregnated with cell-loaded Ag or GelMA or GelMA-Ag, and crosslinked. PCL-hydrogel constructs were then incubated for 4 weeks, and tested for cell viability and collagen production (immunostaining). (B) Coliseum shaped scaffolds were printed, incubated for 2 weeks, and impregnated with cell-loaded GelMA-Ag in the inner region and GelMA in the outer region. After crosslinking, constructs were incubated for 4 weeks and cut into two halves to obtain the meniscus shaped constructs. NS: non-shifted, S: shifted, NSC: non-shifted, with circumferential strands, and SC: shifted and with circumferential strands.

Anatomical constructs were produced using the magnetic resonance imaging (MRI) data of the left knee of a Yucatan miniature pig cadaver scanned on a 3T scanner (Siemens Trio, Erlangen, Germany) as described previously [27]. Briefly, lateral meniscus was manually segmented from both sagittal and coronal planes using the medical image processing software Mimics 9.11 (Materialise, Belgium). A 3D voxel model was reconstructed from the data, converted to STL format and printed.

Scaffolds were sputter-coated with gold and visualized using scanning electron microscope (NanoEye, South Korea). Strand diameter and the distance between strands (pore size) were estimated using ImageJ software (NIH).

### 2.2 Mechanical testing

Cell free PCL scaffolds (n= 4) were incubated in PBS for 24 h and tested under compressive or tensile load using CellScale mechanical tester (Univert, Canada) equipped with a 10 N load cell. Compression and tension (gauge length: 10 mm) testing were performed at a displacement rate of 1 mm.min^−1^ on the square and rectangular prism samples, respectively. Compressive and tensile moduli were calculated from the elastic regions of the stress-strain curves.

### 2.3 Synthesis of GelMA

GelMA was produced as described previously [2,28] with a methacrylation degree of 64%. Briefly, a solution (10% in PBS, pH 7.2) of gelatin (from porcine skin, 300 Bloom, Sigma-Aldrich, USA) was mixed with methacrylic anhydride (MA) (Sigma-Aldrich) in a volume ratio of 5:1 (Gelatin:MA), incubated at 50 °C for 1.5 h, dialyzed against phosphate buffer (0.1 M, pH 7.2) for 3 days, and freeze dried at −80 °C.

### 2.4 Preparation of cell-hydrogel constructs

Human fibrochondrocytes, previously isolated from menisci of a 56 year-old female patient going through a total knee replacement at Hacettepe University Department of Orthopedics (Ankara, Turkey) with her written consent, were a gift from Ars Arthro Biotech Co. (Ankara, Turkey) [13]. Cells were expanded in tissue culture flasks containing culture media (DMEM:F12 (1:1) supplemented with 10% fetal bovine serum (FBS), 100 U/mL penicillin/streptomycin, and 1 μg/mL Amphotericin B (all Thermo Fisher Scientific, USA)) before use.

Cells (passage 3) were suspended in solutions of GelMA (5%, containing 1% (w/v) of the photoinitiator (PI, Irgacure 2959, Sigma-Aldrich)), agarose (Ag, low gelling temperature, Sigma-Aldrich) (2%, kept at 43 °C), and GelMA-Ag (GelMA:Ag, 5:2 w/w) (3.5%, containing 1% PI and kept at 43 °C) in DMEM-F12 media. Cell-polymer suspensions (volume: 100 μL, cell density: 5 × 10^5^ cells/mL media) were placed in polydimethylsiloxane (PDMS)-coated 96-well plates to prevent attachment of the gels on the plate surface after crosslinking. The Ag-containing solutions were cooled down to room temperature, and then the GelMA-containing solutions were exposed to UV radiation (λ: 365 nm) at 0.13 mW/cm^2^ for 3 min to form the cell-loaded hydrogels (Fig. 1A(ii)). Fibrochondrocytes were also seeded on the PCL (SC) scaffolds (cell densities: 5 × 10^4^ cells/scaffold) as the controls. The constructs were incubated for 21 days in culture media. Some samples were removed from culture media at Day 7 and analyzed for cell morphology, while others were analyzed for mRNA expression every week during the 21-day incubation.

### 2.5 Cell morphology

To examine the cell morphology, constructs were fixed with 4% paraformaldehyde, incubated for 5 min at room temperature in Triton X-100 solution (0.1%, v/v in PBS), for 30 min in bovine serum albumin (BSA) solution (1% in PBS), for 30 min in DRAQ5 (red, to stain the nuclei) and for 1 h in Alexa fluor 532-labelled phalloidin (green, to stain the actin) all at 37 °C, and examined using a confocal laser scanning microscope (CLSM) (Leica DM 2500, Germany). The morphology of cells (n≥30) in the CLSM images were analyzed using the ImageJ software, and the area, perimeter, circularity and aspect ratio were determined.

### 2.6 Quantitative real time PCR analysis

For analysis of gene expression, constructs (n=2) were removed from culture at Days 7, 14, and 21, rinsed with PBS, frozen at −80 °C and lyophilized for 10 h. Cells were disrupted using buffer RLT, and the total RNA was extracted using RNeasy Micro Kit (Qiagen, Germany). First strand cDNA was synthesized in a thermal cycler (iCycler, BIO-RAD, USA) using RevertAid™ First Strand cDNA synthesis kit (Thermo Fisher Scientific, USA). SybrGreen Quantitative RT-PCR kit (Sigma Aldrich, Germany) was used with Corbett Rotor-Gene 6000 (Qiagen, Germany) system for qPCR analysis. The sequences of the chondrogenic and fibrochondrogenic gene-specific primers (all purchased from Sentegen, Turkey) used in the assays are presented in Table S1. The results were normalized to GAPDH, and calculated relative to cells incubated on tissue culture polystyrene (TCPS). Relative gene expression was calculated by the ∆∆Ct method. All experiments were performed in duplicates.

### 2.7 Preparation of the PCL-hydrogel constructs

Fibrochondrocytes (passage 3) were seeded on the square prism PCL (SC) scaffolds (cell density: 3 × 10^4^ cells/scaffold), and incubated for 2 weeks in culture media (Fig. 1A(iii)). At Week 2, the square prism scaffolds were impregnated with 50 μL of cell suspensions (cell densities: 6 × 10^5^ cells/mL media) in GelMA (5%, containing 1% of PI) or Ag (2%, kept at 43 °C) or GelMA-Ag (3.5%, containing 1% of PI and kept at 43 °C) (Fig. 1A). Ag-containing constructs were cooled down to room temperature, and then GelMA-containing constructs were exposed to UV for 3 min to form the hydrogels. Thus, PCL (as control), PCL-Ag, PCL-GelMA, and PCL-(GelMA-Ag) constructs were produced. Constructs were incubated for additional 4 weeks in culture media and analyzed at Week 6 for cell viability and collagen production.

The Coliseum shaped PCL scaffolds were similarly seeded with cells (density: 10^6^ cells/scaffold) and incubated for two weeks (Fig. 1B). Scaffolds were impregnated with 0.5 mL of (GelMA-Ag)-cell suspension (cell density: 6 × 10^5^ cells/mL) in the inner region, left to cool and then exposed to UV for 3 min to form the hydrogel. Scaffolds were then impregnated with 1.5 mL of GelMA-cell suspension (cell density: 6 × 10^5^ cells/mL) in the outer region and again exposed to UV for 3 min to crosslink GelMA. Constructs were incubated for 4 weeks in culture media, and cut into two halves at Week 6 to obtain the meniscus shaped constructs (Fig. 1B). Constructs were analyzed for cell viability and collagen production.

### 2.8 Cell viability

Cell viability on the PCL-hydrogel constructs was assessed using Live/Dead™ viability assay kit (Thermo Fisher Scientific). Constructs (n=2) were removed from culture media at Week 6, and stained in a solution containing Calcein-AM (2 μM in PBS) (live cells, green) and ethidium homodimer (EthD)-1 (4 μM) (dead cells, red) for 30 min. Samples were scanned using CLSM down to a depth of about 350 μm (40 images) in z-direction, and z-stack images (*n*=5 images/sample) were reconstructed. Live (green) and dead (red) cell numbers were estimated using ImageJ software and cell viability was calculated.

### 2.9 Collagen deposition

Collagen type I (COL 1) and II (COL 2) deposition was examined using immunostaining. At Day 1 and Week 6, cell containing PCL-hydrogel constructs (n=2) were fixed with paraformaldehyde (4%, w/v) and incubated in Triton X-100 (0.1%, v/v) for 5 min. For double immunostaining, the samples were incubated in mouse primary antibody against human COL 1 (dilution 1:1000), and then in rabbit primary antibody against human COL 2 (both Cell Signaling Technology, USA) (dilution 1:1000) for 1 h each. Samples were then incubated in a mixture of Alexafluor 488-labelled anti-rabbit IgG and Alexafluor 532-labelled anti-mouse IgG (both Cell Signaling Technology) (dilutions: 1:1000) for 1 h at room temperature. Finally, samples were incubated in TO-PRO™-3 iodide (Thermo Fisher Scientific) dye (dilution 1:1000) for 30 min at room temperature. Samples were scanned using a CLSM to thicknesses of 250 μm and 100μm (30 images) in z-direction for the 4x and 10x objectives, respectively. Z-stack images were reconstructed.

COL 1 and COL 2 signal intensities of the z-stack images (n=2-5 images/sample) were quantified using ImageJ. CLSM images of the square prism constructs were converted to 8-bit and the integrated densities (at red (for COL 1) and green (for COL 2) channels) were estimated. To eliminate background fluorescence, Day 1 results were subtracted from Week 6 results. For the meniscus shaped constructs signal intensity was measured along a straight line in radial direction (crossing from the outer region towards the inner region).

### 2.10 Statistical analyses

Statistical analyses were performed using SPSS 23 (IBM, USA). Student’s t-test was performed to compare the morphology of cells on PCL with and without circumferential strands. One-way ANOVA was performed to compare the mechanical properties, cell morphology, quantitative RT-PCR, cell viability and immunohistochemistry results. Two-way ANOVA was performed to test the interaction between two independent variables (presence of shifts and/or circumferential strands for the mechanical tests, and time and material type for the quantitative RT-PCR results). Tukey’s honestly significant difference (HSD) post-hoc tests were performed after ANOVA. Data are presented as the mean ± standard deviation (SD). Significance level *α* was 0.05.

## 3 Results and Discussion

The aim of this study was to engineer a structurally, biochemically, and anatomically correct meniscal construct. We produced a meniscus shaped, 3D printed PCL scaffold and impregnated it with human fibrochondrocyte-loaded GelMA in the outer region and GelMA-Ag in the inner region, to create a bizonal biochemical composition resembling that of the native tissue. Hydrogels are reported to enhance ECM production [2], increase viability of the cells on PCL scaffolds under dynamic load [26], and reduce the damage to the articular cartilage after implantation into cruciate ligament-transected mice [25].

In order to optimize the printing parameters, we first printed PCL scaffolds in various design settings and tested their microarchitecture and mechanical properties.

### 3.1 Effect of scaffold design on microarchitecture and mechanical properties of PCL scaffolds

The PCL scaffolds were designed and printed in square prism shapes, with or without circumferential strands, and with or without shifts (Fig. 2A). Shifts were introduced to decrease the compressive stress exerted on the cartilage, thereby reducing the risk of cartilage degeneration after implantation. Circumferential strands were added to mimic the collagen fibers in the native tissue and reinforce the scaffolds against tensile hook stress. Scaffolds with the circumferential strands (NSC and SC) had more defined shapes than those without the circumferential strands (NS and S); those without the circumferential strands were not perfect square prisms.

**Figure 2.**
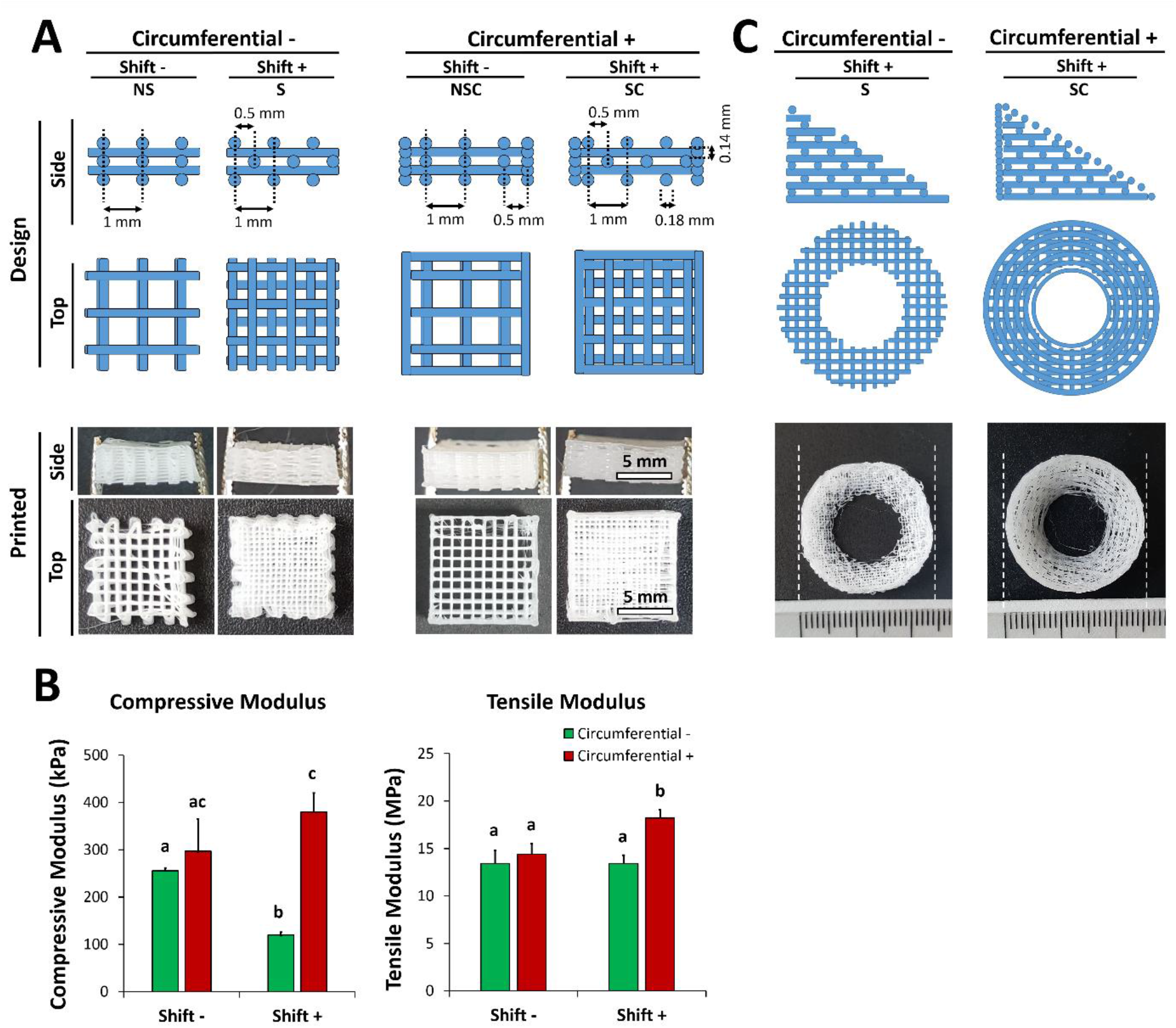
Designs of the 3D printed scaffolds and their effects on mechanical properties. (A) Design and production of the square prism PCL scaffolds. (B) Elastic moduli of the PCL scaffolds under compressive (left) and tensile load (right). n=4. Data are presented as the mean ± SD. (C) Design and production of the meniscus shaped PCL scaffolds.

The average strand diameter was significantly greater (p<0.001) for scaffolds with the shifted interior design (184 ± 6 μm) than that of scaffolds with the non-shifted design (163 ± 10 μm), probably because the strands in the shifted design widened during printing because they sat on the underlying strands (Fig. S1). The distance between strands (pore size) did not change with shifting. Pore size was 810 ± 40 μm for all constructs, large enough to allow for hydrogel infiltration into the scaffold center and easy transport of nutrients and waste products. The optimal pore size range is between 150 and 500 μm for meniscal tissue infiltration [29], and around 200 μm for cell proliferation and ECM production [18]. The pores of our scaffolds are larger than the optimal values; however, impregnation of our scaffolds with hydrogels would reduce the pore size to the desired range.

In wet state, the scaffolds exhibited compressive moduli in the range 150-400 kPa, and tensile moduli in the range 13-18 MPa (Fig. 2B). Shifting of strands alone resulted in reduced compressive modulus (from 256 ± 6 kPa for (NS) to 120 ± 6 kPa for (S)), but had no effect on tensile modulus (from 13.4 ± 1.4 MPa to 13.4 ± 0.9 MPa). On the other hand, when circumferential strands were introduced alone (NSC), both compressive (297 ± 68 kPa) and tensile (14.4 ± 1.1 MPa) moduli increased. The highest compressive (380 ± 40 kPa) and tensile (18.2 ± 0.9 MPa) moduli (p<0.05) were obtained when both shifted and circumferential strands (SC) were present (the two factors had interaction, two-way ANOVA, p<0.01). The circumferential strands in the exterior of the scaffolds probably fused with the adjacent shifted strands in the interior, and formed a more condensed and rigid overall structure (Fig. 2A).

Compressive modulus of the PCL (SC) scaffolds was within the modulus range of the native human meniscus (0.3-2 MPa) [30,31]. Tensile modulus was comparable to that of native human meniscus calculated in radial direction (4-20 MPa), but not in circumferential direction (78-125 MPa) [32].

The 3D printed PCL scaffolds prepared by other researchers for meniscal regeneration had higher tensile (30-80 MPa) and compressive (10-54 MPa) moduli than ours, probably because they printed their scaffolds with larger strand diameter (0.3 mm) and smaller strand distance (0.1-1.0 mm) [18–22] than our scaffolds (strand diameter: 0.18 mm, strand distance: 1 mm). However, their scaffolds were reported to degenerate the cartilage after implantation, due to high stiffness [20,21]. Although it is possible to increase the mechanical properties of our constructs by changing the strand diameter and/or strand distance, we preferred to keep the compressive modulus in the lower range of the native meniscus in order to reduce degeneration of the cartilage surface. In fact, in our previous studies we used a high molecular weight (MW) PCL (80 kDa) to print our constructs, and the compressive and tensile moduli of the constructs were 10 and 30 MPa, respectively [26]. In the current study, we used a lower MW PCL (50 kDa) and printed our constructs with shifted design, which resulted in lower compressive properties (modulus 400 kPa) and thus reduced the risk of cartilage degeneration in the expense of lowered tensile modulus (18 MPa). It is known, however, that the mechanical properties of PCL scaffolds increase upon seeding with cells or after implantation [10,21,33]. So, it is expected that the mechanical properties of our constructs would get closer to those of the meniscus after possible implantation into patients’ knees.

### 3.2 Cell morphology and gene expression on PCL scaffolds and hydrogels

PCL (SC) scaffolds exhibited the highest mechanical properties –the closest to meniscus. Therefore, this design was chosen for use in the rest of the study. Coliseum shaped scaffolds printed with the circumferential strands and shifted design (SC) had more defined shape than those with shifted design but without the circumferential strands (S) (Fig. 2C).

We examined the effect of circumferential strands on cell morphology. Fibrochondrocytes were incubated for 7 days on PCL (S and SC) scaffolds, and on Ag, GelMA, and GelMA-Ag hydrogels. Cells were elongated and aligned along the circumferential strands, but randomly distributed and more elongated when no circumferential strands were present, especially at the contact points where the two PCL strands conjoined (Fig. 3A). Cells on GelMA were spread and elongated without a certain alignment pattern, while those on Ag were round. On GelMA-Ag, some of the cells were round and some were spread/elongated. Semi-quantitative analysis using ImageJ revealed that cell area and circularity significantly decreased (p<0.0001), while aspect ratio significantly increased (p<0.0001) upon introduction of circumferential strands (Fig. 3B). Cell perimeter, on the other hand, did not change with the presence of circumferential strands. These results clearly showed that circumferential strands induced cell elongation and alignment. Alignment of cells is of great importance for secretion of aligned collagen fibers, which would contribute to tensile strength of the tissue [3,4]. The reason for the elongated cell morphology on PCL strands could be the stiffness of this material. Fibrochondrocytes were reported to get more elongated with the increasing substrate stiffness [9]. PCL scaffolds could also force the cells to elongate, because they are produced in the form of strands (1D), which have a high aspect ratio [34].

**Figure 3.**
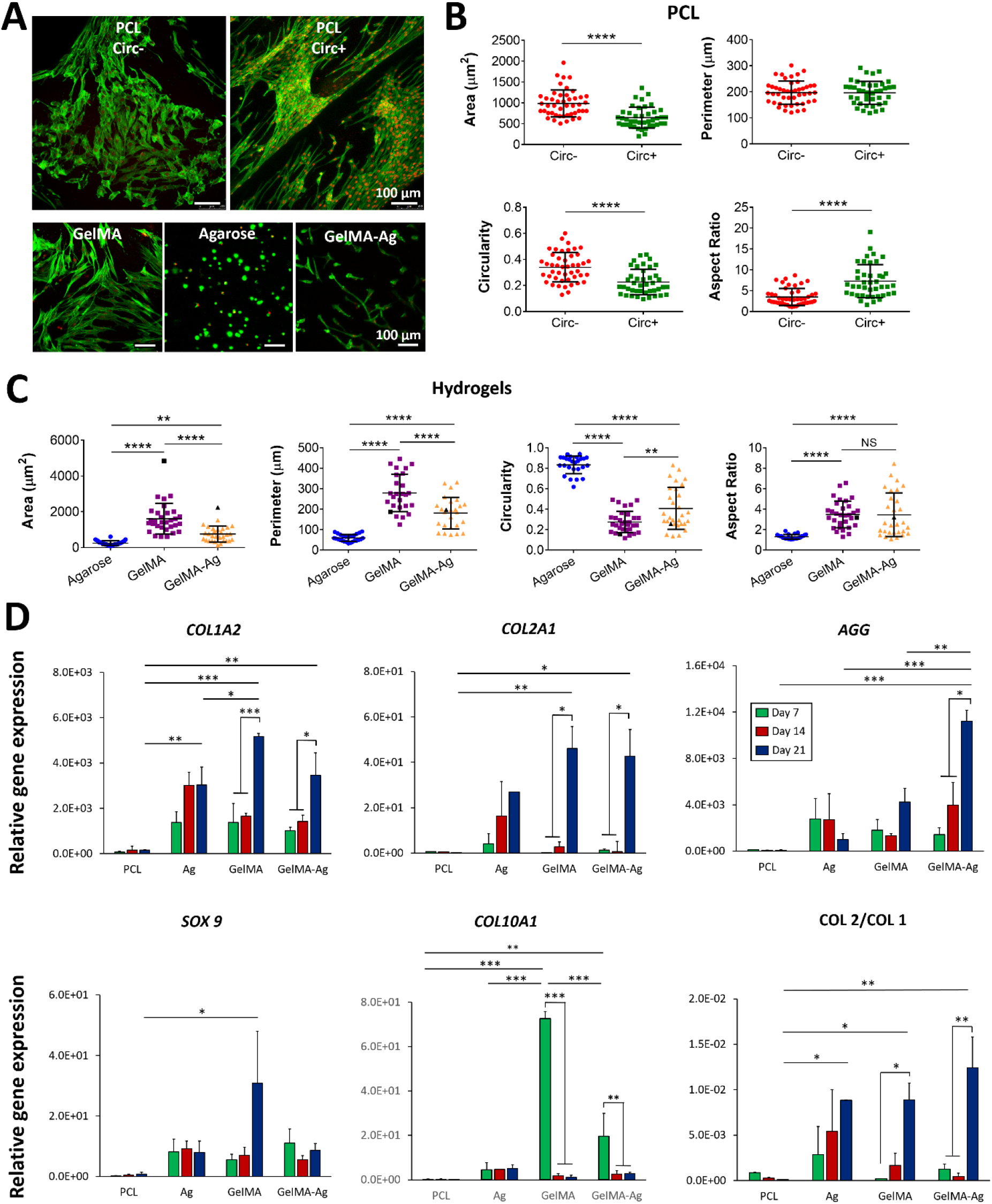
Cell morhology and gene expression on the constructs. (A) Confocal laser scanning microscopy (CLSM) images showing cell morphology on the surface of the meniscus-shaped PCL scaffolds and the hydrogels after 7 days of incubation. Nuclei: red (DRAQ5), and actin: green (Alexa fluor 532-labelled phalloidin). (B, C) Semi-quantitative analysis of the CLSM images in (A) showing cell area, perimeter, circularity and aspect ratio on (B) the PCL without (circ-) and with (circ+) circumferential strands, and on (C) the hydrogels. (D) Quantitative real time PCR showing gene expression on the materials relative to that on TCPS. *COL1A2*: Type I collagen, *COL2A1*: type II collagen, *AGG*: aggrecan, *SOX 9*: SRY-box 9, and *COL10A1*: type + collagen. All results are normalized to GAPDH. Data are presented as the mean ± SD. n=2. *p<0.05, **p<0.01, ***p<0.001, and ****p<0.0001.

On hydrogels, the greatest cell area and perimeter were on GelMA (p<0.0001), and the smallest on Ag (Fig. 3C). Circularity, on the other hand, was the smallest on GelMA (p<0.01) and the greatest on Ag (p<0.0001). Introduction of GelMA to Ag resulted in reduced circularity. On the other hand, cell aspect ratio was the smallest in Ag (p<0.0001), while cells in GelMA and GelMA-Ag exhibited similar aspect ratio values. Similar observations were reported in our previous study, in which we showed that cells interact more strongly with GelMA leading to a more spread morphology [2]. Gelatin has biologic recognition sites such as arginine-glycine-aspartic acid (RGD) sequences which promote cell adhesion [35]. On the other hand, Ag lacks such biologic recognition sites [36] and is highly hydrophilic, which leads to low spreading of the cells on Ag hydrogels.

The different cell-material interactions not only affected cell morphology, but also led to different gene expression profiles. Hydrogels exhibited higher levels of gene expression than PCL (SC) scaffolds, and the level of gene expression generally increased over time (Fig. 3D). GelMA exhibited high levels of *COL1A2*, *COL2A1, SOX 9*, and *COL10A1* expression after 21 days of incubation, while GelMA-Ag exhibited high levels of *COL2A1* and *AGG* expression. The RGD sequences on gelatin [35] probably induced expression of *COL1A2* (300-400 fold higher on GelMA than PCL (p<0.001)) and *COL2A1* (100-200 fold higher on the GelMA-containing gels than PCL (p<0.01)). In line with our results, chondrocytes were reported to exhibit higher levels of *COL1A2* and *COL2A1* on GelMA than on the PCL-reinforced GelMA after 14 days of incubation [37]. The high level of *SOX9* on GelMA is also expected, since *SOX9* is known to support expression of type II collagen [38]. *COL10A1* is generally associated with late stage chondrocyte hypertrophy and/or endochondral matrix remodeling, and is expressed in mesenchymal stem cells (MSCs) going through chondrogenesis after 21 days of incubation [39]. In the present case, *COL10A1* expression was the highest on GelMA hydrogels and it decreased after Day 7. This indicates that fibrochondrocytes were remodeling the matrix in the first week of incubation probably by degrading GelMA. This also explains the low levels of *COL1A2* and *COL2A1* on the GelMA-containing gels during the first two weeks of incubation. Instead of producing new collagen (type I and II), cells probably remodeled the existing collagen (which came from GelMA). In fact, type × collagen was also reported to be expressed during development of healthy fibrochondrocytes [40] and even in MSCs [41], which showed that it should not necessarily be associated with hypertrophy.

On the other hand, PCL and Ag lack RGD sequences, and the main parameters regulating gene expression on these materials are their chemistry and stiffness. PCL is hydrophobic, which limits initial cell attachment. It is also stiff, which directs cells to proliferate rather than produce ECM [2]. This explains the lower levels of *COL1A2* (p<0.001) and *COL2A1* expression on PCL than on the hydrogels (Fig. 3D). Conversely, Ag lacks RGD sequences and its gel is soft and highly hydrated. Thus, cells take up a round morphology in Ag gels, which contributes to cartilaginous gene expression [2,26]. In fact, *COL1A2* and *COL2A1* expression on Ag was high during the first two weeks of incubation. The low level of *COL10A1* on Ag also shows that no matrix remodeling was required on this gel, which might explain the comparable levels of *COL1A2* and *COL2A1* on Ag with (or even higher than) those on the GelMA-containing gels in the first two weeks of incubation.

Interestingly, GelMA-Ag exhibited the highest *AGG* expression after 21 days of incubation (Fig. 3D). When preparing GelMA-Ag hydrogels, first Ag was set and then GelMA was UV crosslinked. This created Ag-rich domains within the gel, which prevented the formation of densely packed GelMA fibers. The high expression of *AGG* on GelMA-Ag gels could be a result of this heterogeneity. Heterogenic domains are reported to contribute to growth and maturation of the fibrocartilage [33] and cartilage [42]. In fact, in our study, both round and elongated cells were present on GelMA-Ag hydrogels (Fig. 3A), suggesting the presence of heterogenic domains in these gels. Besides, *COL10A1* expression was lower on GelMA-Ag than on GelMA gels. Since Ag disrupts packing of the GelMA fibers, there would be less need for matrix remodeling in GelMA-Ag gels. In fact, strain stiffening and remodeling properties of collagen was shown to be disrupted in the presence of Ag in collagen-Ag gels [43].

Finally, *COL2A1*/*COL1A2* ratio, the chondrogenic differentiation index [44], was higher on Ag than the GelMA-containing gels especially in the first two weeks of incubation, but then it increased to a higher level on GelMA-Ag on Day 21 (Fig. 3D). This suggests that the Ag-containing gels are chondrogenic and appropriate for use in the inner region of the constructs, which is known to be cartilaginous [1]. With the highest *AGG* expression being on GelMA-Ag, we chose to use this gel at the inner portion of the constructs.

### 3.3 Effects of hydrogel incorporation on cell viability and ECM production

After examining the effects of PCL and hydrogels on cell morphology and gene expression, we impregnated the cell seeded PCL scaffolds with the cell-loaded Ag, GelMA, and GelMA-Ag to obtain the PCL-Ag, PCL-GelMA, and PCL-(GelMA-Ag) constructs, respectively, and investigated the cell viability and collagen production on these constructs. Cell viability after 6 weeks of incubation ranged from 80% to 93%, with no significant difference between construct types (Fig. 4A). Addition of hydrogels did not have a significant effect on cell viability. Moreover, cell viability in the constructs with the non-shifted PCL scaffolds (Fig. 4A) did not differ from constructs with the shifted design (Fig. 4B). Our results are in line with the previous studies reporting around 80% cell viability on the PCL-Ag and PCL-GelMA constructs at Day 1 [23], Day 7 [37] or Day 56 [26] of incubation after printing.

**Figure 4.**
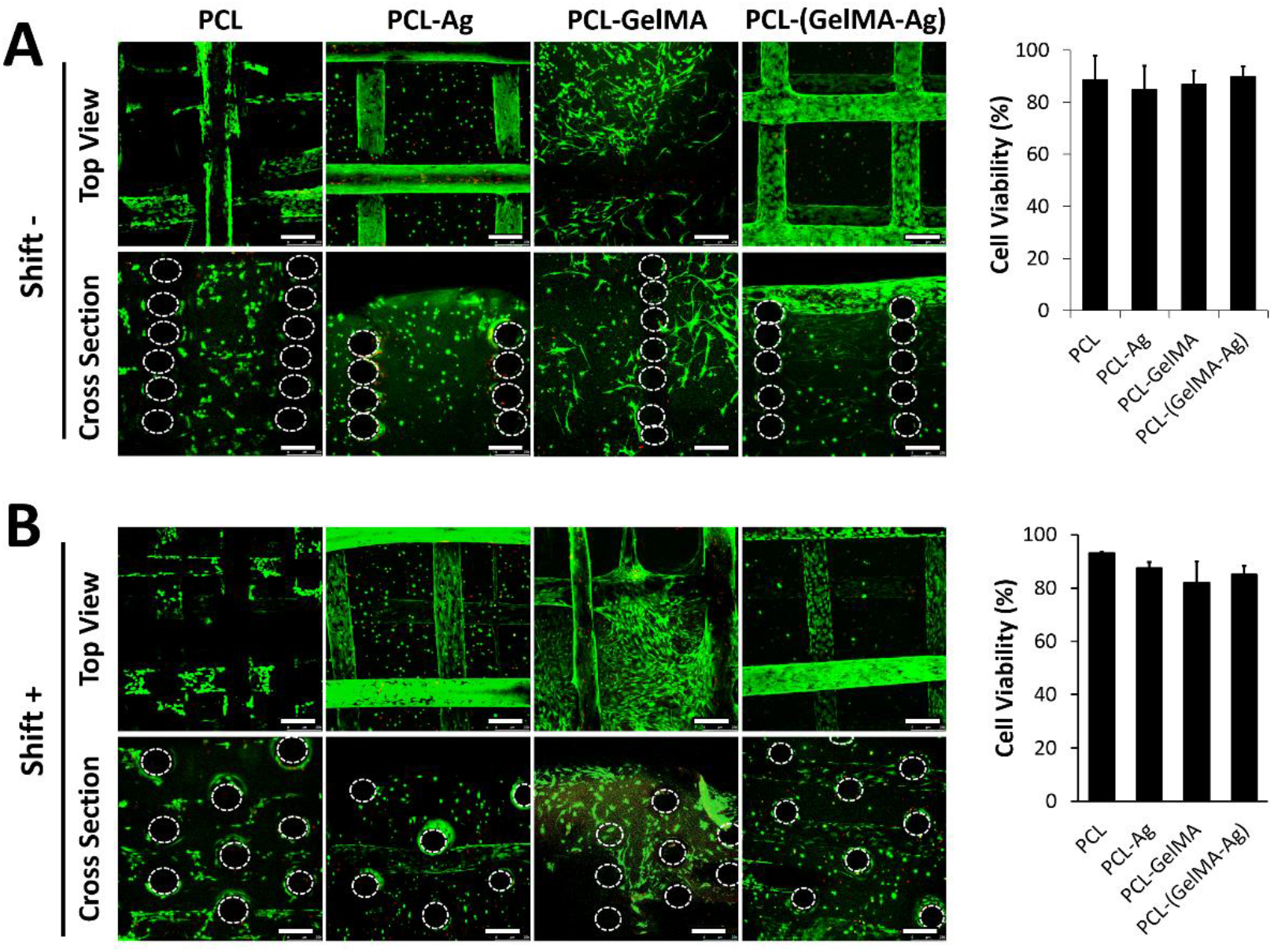
Cell viability on the constructs after 6-week incubation. CLSM images of the constructs with (A) non-shifted, and (B) shifted design after live/dead cell viability assay (left), and semi-quantitative analysis of the CLSM images (n=5 images/sample) showing cell viability on the constructs (right). n=2. Live cells: green (calcein-AM); and dead cells: red (ethidium homodimer-1, EthD-1). Dashed circles: PCL strands orthogonal to view plane. Scale bar: 250 μm.

Immunostaining of the whole PCL-hydrogel constructs revealed low collagen deposition on PCL (Fig. 5A) and PCL-Ag (Fig. 5B), and high deposition on PCL-GelMA (Fig. 5C) and PCL-(GelMA-Ag) (Fig. 5D). GelMA exhibited some background autofluorescence, which obscured the signal coming from the collagen produced by cells (Fig. 5C). Collagen deposition increased on all the constructs after 6 weeks of incubation (Week 6 compared to Day 1 results), and in general COL 1 staining was more intense than COL 2. Similar results were obtained with constructs having the non-shifted design (Fig. S2).

**Figure 5.**
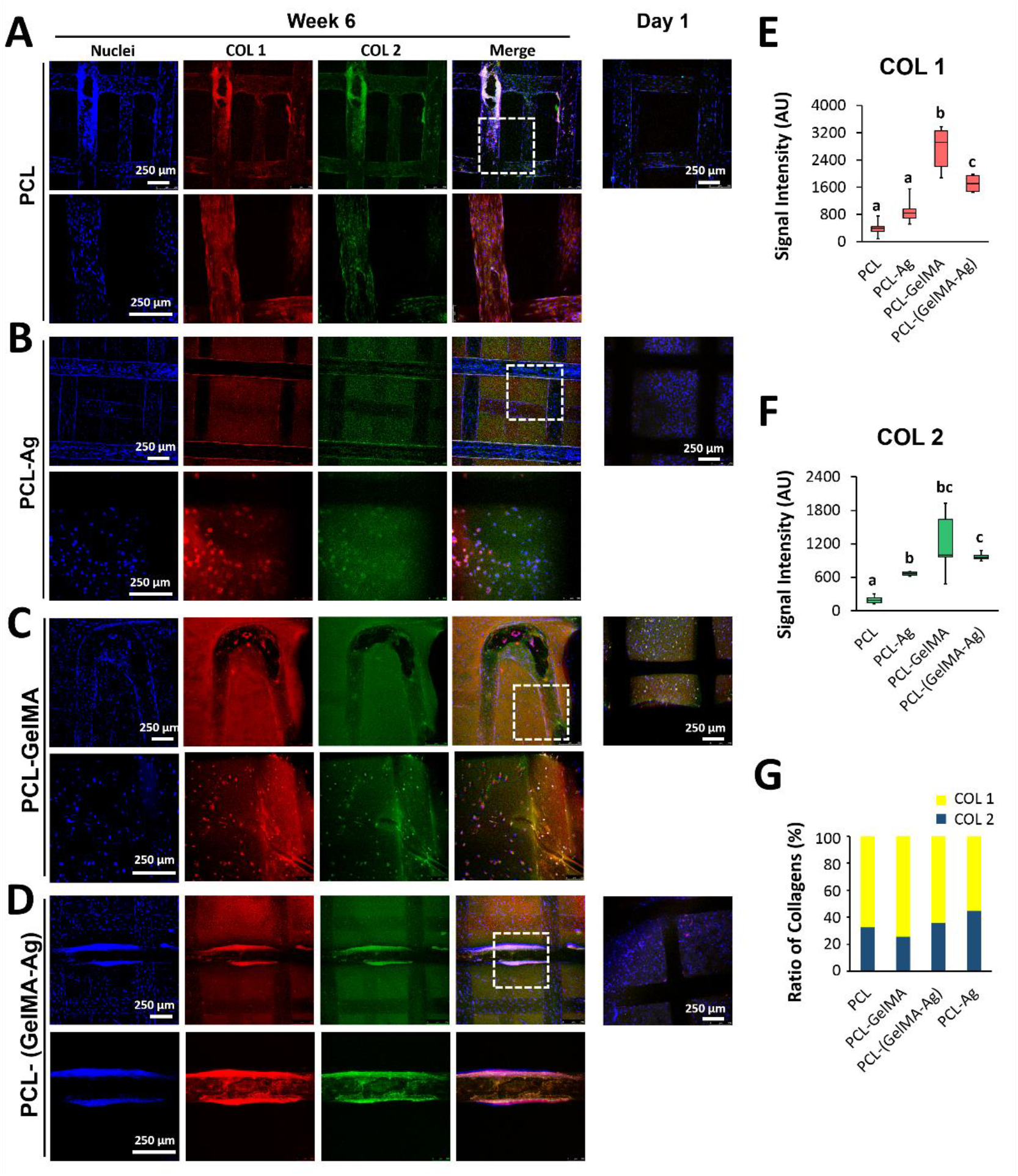
Immunostaining showing deposition of type I (COL 1) and type II (COL 2) collagen on the square prism constructs after 6-week incubation. CLSM images of (A) PCL, (B) PCL-Ag, (C) PCL-GelMA, and (D) PCL-(GelMA-Ag) constructs. Semi-quantitative analysis of the CLSM images showing the intensity of (E) COL 1, and (F) COL 2 signal intensities, and (G) percentage of COL 1 and COL 2. n=2-5. Nuclei: blue (TO-PRO); COL 1: red (Alexa fluor 532 labelled antibody), COL 2: green (Alexa fluor 488 labelled antibody). Dashed squares: magnified regions. Data are presented as the mean ± SD. ^a,b,c,d^Significant difference between results when no letters in common.

Semi-quantitative analysis, which was done after reducing the background autofluorescence (by subtracting Day 1 results from Week 6), revealed that the highest intensity of COL 1 was on PCL-GelMA (p<0.05) (Fig. 5E). Addition of GelMA to PCL enhanced COL 1 deposition by 10 folds (p<0.01), and the GelMA-containing constructs exhibited significantly higher COL 1 staining than PCL and PCL-Ag (p<0.05). The highest COL 2 deposition, on the other hand, was on PCL-GelMA and PCL-(GelMA-Agboth exhibiting similar COL 2 staining (Fig. 5F). Addition of hydrogels to PCL also increased COL 2 deposition significantly (p<0.05).

On PCL, COL 2 constituted around 30% of the total collagen produced (COL 1 and COL 2), and GelMA incorporation decreased this value to around 25% while Ag incorporation increased it to around 45% (Fig. 5G). For PCL-(GelMA-Ag), COL 2 was around 35% of the total collagen. These data are consistent with the gene expression results, which revealed high expression of *COL1A2* and *COL2A1* on the GelMA-containing gels, and a high *COL2A1*/*COL1A2* ratio on the Ag-containing gels (Fig. 3D).

Others also reported high COL 1 production in GelMA gels and high COL 2 production in Ag gels [23], supporting our findings. Similarly, increased COL 2 production was reported on the PLA scaffolds after impregnating them in alginate hydrogel [12]. These results could be attributed to the cell-material interactions and cell morphology, as mentioned earlier. In fact, nucleus pulposus cells embedded in GelMA were reported to assume more rounded morphology and exhibit enhanced chondrogenesis after inhibition of focal adhesion kinases (FAKs) on these cells [45].

### 3.4 Production of structurally, biochemically and anatomically correct meniscus constructs

Since we found that GelMA induced fibrous phenotype (high expression of type I collagen at gene and protein levels), and GelMA-Ag induced cartilaginous phenotype (high aggrecan expression and COL 2/COL 1 index), we impregnated the inner region of the fibrochondrocyte-seeded meniscus-shaped PCL (SC) scaffolds with the cell-loaded GelMA-Ag and the outer region with cell-loaded GelMA at Week 2 of incubation to produce the PCL-dual hydrogel constructs, and further incubated the constructs for additional 4 weeks in culture media. The reason for pre-seeding of PCL was to give cells enough time to deposit ECM on the PCL strands so that the cell-hydrogels attach better on the strands. The hydrogels integrated well with the PCL scaffold (top view) and with each other (cross sectional view) (Fig. 6A), and they were retained for 6 weeks in culture media without any sign of disintegration (Fig. 6B). Live/dead assay also revealed high cell viability on the constructs after 6 weeks of incubation, and no difference was found between cell viabilities of the inner and outer regions of the constructs (both around 80%) (Fig. 6C).

**Figure 6.**
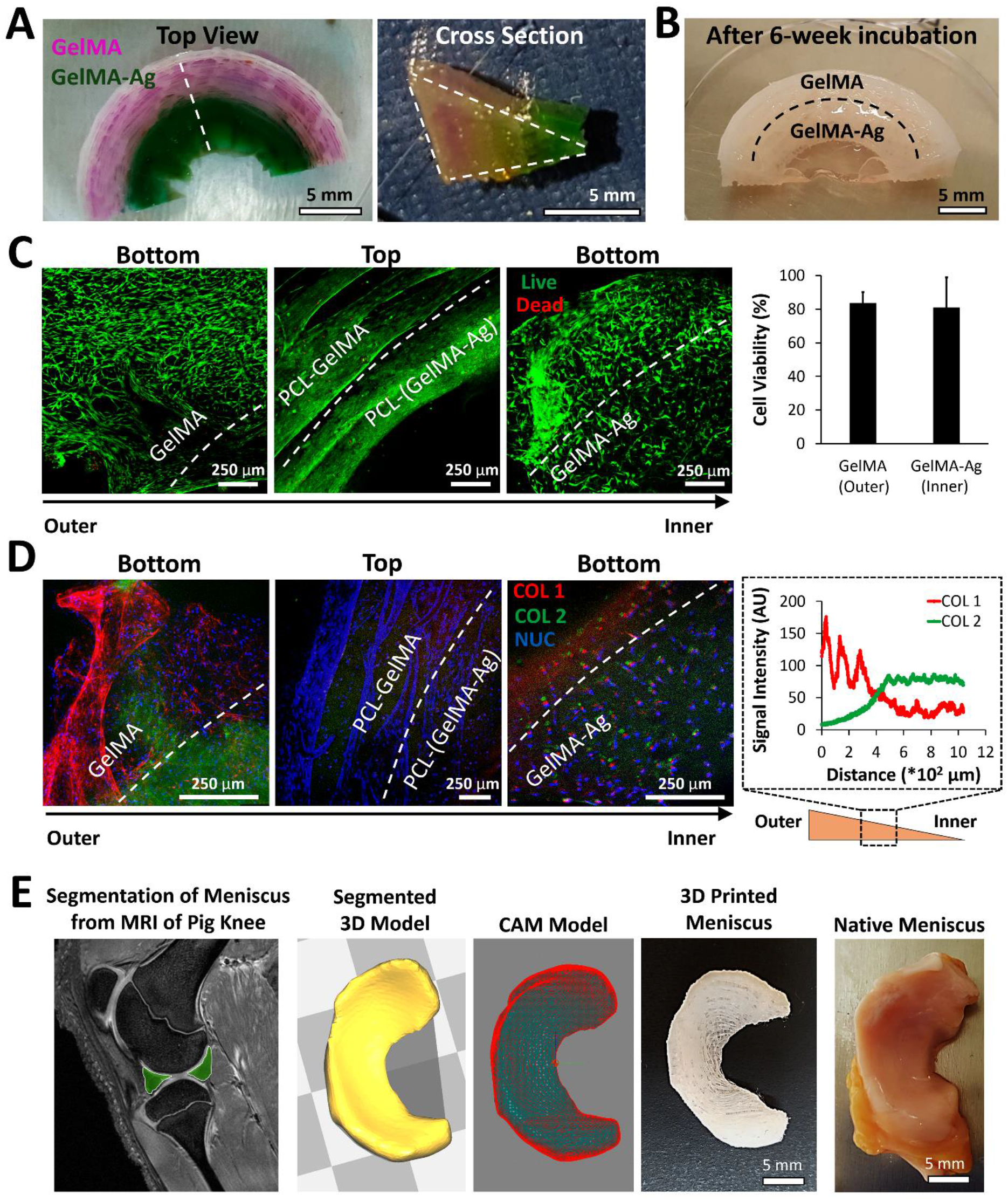
Engineered constructs resembling the biochemical composition, structural organization and anatomy of the native meniscus. (A) The 3D printed PCL scaffold was impregnated with GelMA (phenol red) in the outer region, and with GelMA-Ag (bromocresol green) in the inner region. Top (left) and cross sectional (middle) views.(B) Fibrochondrocyte-seeded construct after 6 weeks of incubation. (C) Cell viability on the constructs after 6-week incubation. CLSM images after live/dead staining (left). Live cells: green (calcein-AM); and dead cells: red (ethidium homodimer-1, EthD-1). Semi-quantitative analysis of the images showing cell viability on different regions of constructs (right). Data are presented as the mean ± SD. n=3. (D) Immunostaining showing collagen deposition on the constructs after a 6-week incubation. CLSM images (left). Nuclei (NUC): blue (TO-PRO); COL 1: red (Alexa fluor 532 labelled antibody), COL 2: green (Alexa fluor 488 labelled antibody). Quantification of the collagen signal intensity along a line crossing from the outer region towards the inner region of the constructs (right). (E) Anatomically correct scaffold was printed after segmentation of the lateral meniscus from the MRI data of a miniature pig knee. CAM: computer aided manufacturing.

The constructs exhibited a high level of COL 1 in the outer region and a high level of COL 2 in the inner region (Fig. 6D). Semi-quantitative analysis showed that the intensity of COL 1 decreased from the outer region towards the inner region, whereas the intensity of COL 2 increased. The COL 2/COL 1 ratio was 10-20% in the outer region and increased to 60-80% towards the inner region. This biochemical composition is very close to that of the native meniscus, at which COL 1 constitutes 80% of the outer region and COL 2 constitutes 60% of the inner region [2].

Large animal models are essential for the preclinical assessment of tissue engineered menisci [46]. The anatomical fit of the implant is decisive as it affects the knee mechanics directly. For a clinically relevant porcine model, we produced anatomically shaped scaffolds using MRI data of a miniature pig knee (Fig. 6E). The PCL scaffolds were printed with SC design, which involved the circumferentially oriented strands mimicking those in the native meniscus. The construct was similar in shape to the native porcine meniscus. This showed that production of patient-specific constructs was possible.

Few studies took the structural organization and biochemical composition into consideration when designing meniscal scaffolds. In one study, electrospun PCL scaffolds were coated on one side with extracts from the digest of the inner meniscus, and on the other side with extracts from the outer meniscus [11]. However, electrospun fiber mats are too thin to form 3D constructs, and have small gaps between the fibers restricting cell migration and infiltration to the scaffold center [10]. In the other study, biochemical composition of the meniscus was mimicked by a cryostructuring method used on lyophilized foams [8], but the foams exhibited low mechanical properties. Finally, 3D printed PCL was tethered with microspheres carrying TGF-β1 in the inner region and those carrying CTGF in the outer region [20]. The scaffold exhibited good mechanical properties; however, the scaffolds led to degeneration of the hyaline cartilage after implantation into a sheep model [20,21]. So, there is a need for reliable scaffolds that closely mimic the structure and biochemical composition of the meniscus, but do not degenerate the cartilage. The advantage of our construct is that it uses Ag and GelMA hydrogels, which are known to reduce the risk of degeneration [25,26]. Our construct would probably fill that gap, and could be used in total replacement of the meniscus.

## 4 Conclusions

In this study, our goal was to produce a meniscal construct that closely mimics the native tissue. To this end, we have printed PCL scaffolds with shifted or non-shifted designs and with or without circumferential strands. We have shown that circumferential strands lead to elongation and alignment of the fibrochondrocytes, which is key to produce circumferentially aligned collagen fibers. We have also shown that hydrogels enhance the ECM production both at mRNA and protein levels; GelMA is fibrogenic (induces expression of COL 1), and GelMA-Ag is chondrogenic (induces expression of aggrecan and yields a high ratio of COL 2/COL 1). Thus, we impregnated the outer region of the meniscus shaped PCL scaffolds with GelMA and the inner region with GelMA-Ag. We have created a construct with increasing COL 1 and decreasing COL 2 expression towards the outer region. This is similar to the native meniscus, the inner portion of which is cartilage-like and the outer portion is fibrocartilage-like. We have produced anatomically correct constructs using the MRI data of a native miniature pig meniscus. Thus, we have successfully created a structurally, biochemically and anatomically correct meniscal constructs. Our construct would be advantageous over the previously proposed PCL scaffolds in that it contains hydrogels which we believe would protect the articular cartilage from degeneration resulting from friction between the stiff PCL scaffolds and the hyaline cartilage. Future studies will involve co-printing of the hydrogels with the PCL to produce a more translational approach, and testing of our construct in a miniature pig model to evaluate its functionality *in vivo*. Our design is valuable in engineering not only the meniscus, but also other tissues that are anisotropic and heterogenic in nature.

## Acknowledgements

This project was supported by BIOMATEN, METU [grant number BAP-08-11-2016-022]; the Ministry of Industry and Commerce of Turkey (SanTez 00356.STZ.2009-1); and the U.S. Department of Veterans Affairs Rehabilitation Research and Development Service [grant number 5IK2RX000760]. GB was supported by TUBITAK [through Bideb 2214/A]. Ars Arthro Biotech. Co. (Ankara, Turkey) is acknowledged for providing the human fibrochondrocytes. Dr. Braden C. Fleming, PhD is acknowledged for kindly sharing the MRI data. The authors declare that they have no competing interests. All the data needed to evaluate the conclusions made in this paper are present within the data presented in the paper and/or the Supplemental Materials. Additional data may be requested from the authors.

## Data Availability

The raw data required to reproduce these findings will be provided upon request.

