## Supplemental Material for "Anatomical meniscus construct with zone specific biochemical composition and structural organization" (from another version of this preprint: version 1, 2019-06-10)

**Supplemental Data**

**Table S1.** Primer sequences for qPCR.

| **Marker** | | **Primer (5' → 3')** | **Amplicon Size (bp)** |
|---|---|---|---|
| GAPDH | Forward | AATCCCATCACCATCTTCC | 155 |
| | Reverse | GCAGAGATGATGACCCTTT | |
| COL1A2 | Forward | CCTAGCAACATGCCAATCTTTAC | 181 |
| | Reverse | AGCAAAGTTCCCACCGAGAC | |
| COL2A1 | Forward | GAGAAGGGAGAGCCTGGAGA | 118 |
| | Reverse | ATCCTCTCTCACCACGTTGC | |
| AGG | Forward | AGGGCGAGTGGAATGATGTT | 79 |
| | Reverse | CGTTTGTAGGTGGTGGCTGT | |
| SOX9 | Forward | ACTCGCCACACTCCTCCTC | 138 |
| | Reverse | CCTCTCGCTTCAGGTCAGC | |
| COL10A1 | Forward | AGAATCCATCTGAGAATATGC | 187 |
| | Reverse | CCTCTTACTGCTATACCTTTAC | |

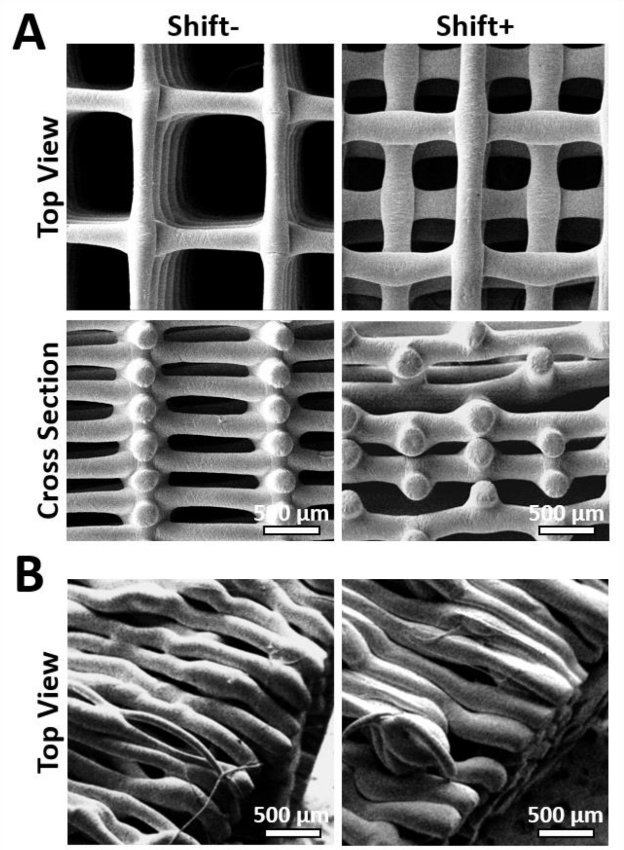


**Figure S1.** Scanning electron microscographs of the 3D printed PCL scaffolds. (A) Square prism, and (B) meniscus shaped scaffolds with non-shifted (Shift-) and shifted (Shift+) designs.

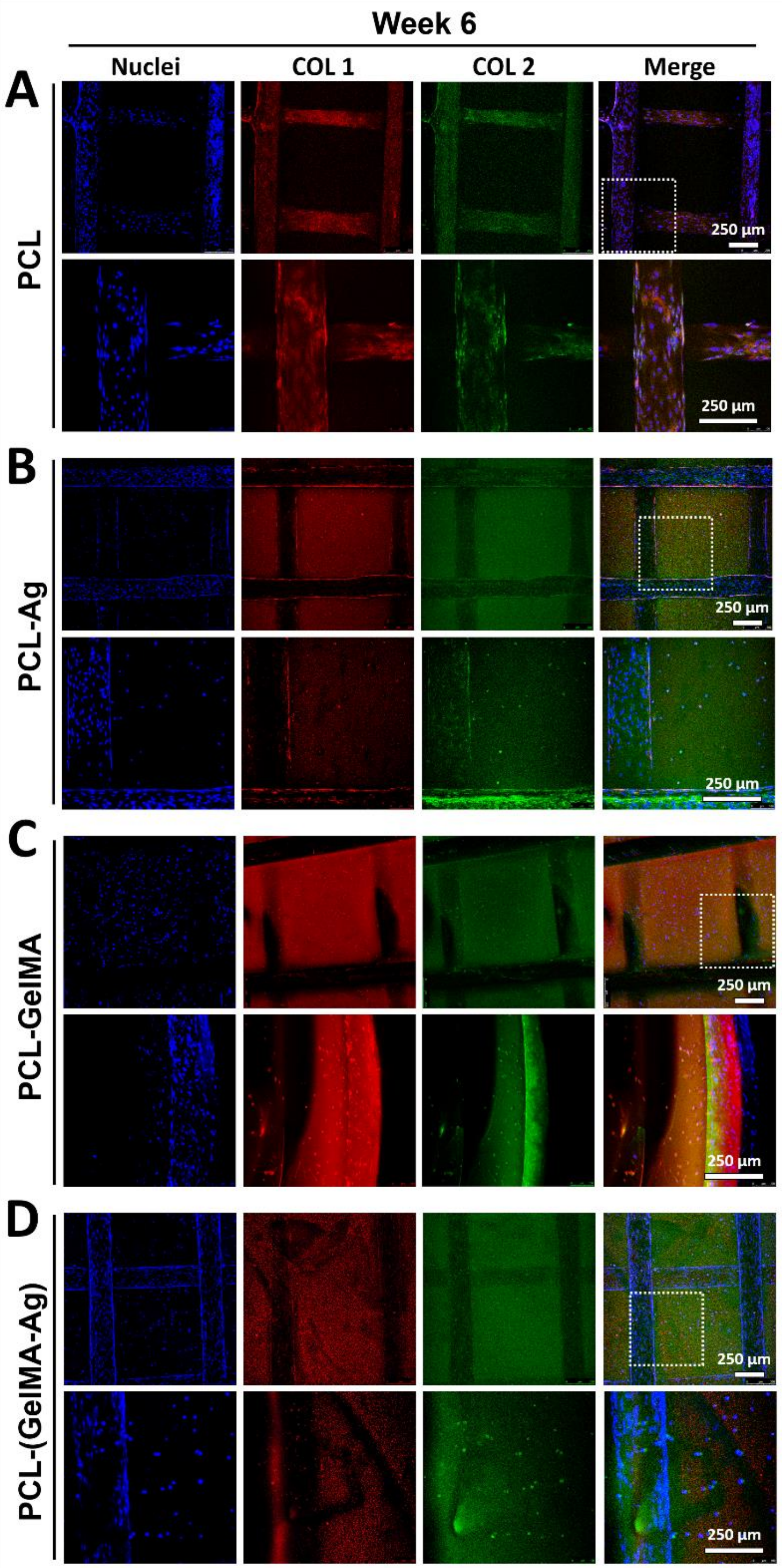
Week 6
Nuclei
COL 1
COL 2
Merge
A
PCL
250 µm
250 µm
B
PCL-Ag
250 µm
250 µm
C
PCL-GelMA
250 µm
250 µm
D
PCL-(GelMA-Ag)
250 µm
250 µm

**Figure S2.** Immunostaining showing deposition of type I (COL 1) and type II (COL 2) collagen on the constructs with non-shifted design after a 6-week incubation. (A) PCL. (B) PCL-Ag, (C) PCL-GelMA, and (D) PCL-(GelMA-Ag). Nuclei: blue (TO-PRO); COL 1: red (Alexa fluor 532 labelled antibody), COL 2: green (Alexa fluor 488 labelled antibody). Dotted squares: magnified regions.
